# Human es-fMRI Resource: Concurrent deep-brain stimulation and whole-brain functional MRI

**DOI:** 10.1101/2020.05.18.102657

**Authors:** WH Thompson, R Nair, H Oya, O Esteban, JM Shine, CI Petkov, RA Poldrack, M Howard, R Adolphs

## Abstract

Mapping the causal effects of one brain region on another (effective connectivity) is a challenging problem in neuroscience, since it requires invasive direct manipulation of brain function, together with whole-brain measurement of the effects produced. Here we establish a unique resource and present data from 26 human patients who underwent electrical stimulation during functional magnetic resonance imaging (es-fMRI). The patients had medically refractory epilepsy requiring surgically implanted intracranial electrodes in cortical and subcortical locations. One or multiple contacts on these electrodes were stimulated while simultaneously recording BOLD-fMRI activity in a block design. Multiple runs exist for patients with different stimulation sites. We describe the resource, data collection process, preprocessing using the fMRIPrep analysis pipeline and management of artifacts, and provide end-user analyses to visualize distal brain activation produced by site-specific electrical stimulation. The data are organized according to the brain imaging data structure (BIDS) specification, and are available for analysis or future dataset contributions on openneuro.org including both raw and preprocessed data.

## Background and Summary

Direct manipulation of brain function is typically considered the gold standard to establish causal relations among brain networks and behavior, and is now widely achieved with techniques such as optogenetics in animals 1. However, direct manipulation of brain function in humans is inherently difficult. Noninvasive methods, such as TMS or tDCS, lack the spatial resolution required for circuit-level specificity. Invasive methods, such as deep brain stimulation (DBS), have been applied for some time, and quite successfully in the case of movement disorders. In addition, DBS is being used to treat a number of other disorders, and is used in an acute setting to help guide neurosurgical decisions for the treatment of epilepsy.

Direct electrical stimulation of the human brain has an extensive literature 2,3 documenting profound acute effects resulting from stimulation of a variety of structures on perception 4, cognition 5, and emotion 6. These findings suggest that the positive benefits of DBS stem from the modulation of systems-level organization in the brain. Consistent with this notion, alterations in brain connectivity are now thought to underlie much of psychopathology 77,89. Both invasive 9,10 and noninvasive 11 neurostimulation are regularly used, and gaining popularity, to treat a number of neurological and psychiatric diseases, including depression 9,11 and memory disorders 12. For instance, deep-brain stimulation of the subgenual prefrontal cortex was discovered as a novel treatment for medically refractory depression 13 and is continuing to be explored 14. However, there is currently a relatively poor understanding of the precise effects, stimulation locations and protocols required to appropriately facilitate clinical improvements in complex, neuropsychiatric syndromes. A major reason underlying this challenge is that focal brain stimulation does not produce cognitive or behavioral effects directly -- it has to act through the rest of the brain. A fundamental problem in systems and cognitive neuroscience is therefore to visualize and understand how intervention at one specific node (stimulation of a specific brain region) influences function everywhere else in the brain.

One solution to this problem is to combine brain stimulation with whole-brain neuroimaging, making it possible to quantify long-range and network-level causal effects produced acutely by the stimulation. While the combination of these techniques is possible in nonhuman primates as animal models, in humans it is typically impossible due to engineering and safety challenges. We have developed a protocol in patients who are undergoing invasive epilepsy monitoring for concurrent electrical stimulation and functional magnetic resonance imaging (es-fMRI) at 3 Tesla. This novel technique in humans offers concurrent invasive electrical stimulation and fMRI-BOLD (blood-oxygen-level-dependent) recordings in the same subject 15, enabling the investigation of new questions about whole-brain activity in relation to focal modulation of brain function. The stimulated regions can include both cortical and subcortical regions, and can include stimulation through depth electrodes or through cortical grids, depending on where electrodes have been clinically implanted in order to monitor seizures. es-fMRI produces reliable distal activations with stimulation protocols of about 10 minutes 15,16.

In this manuscript, we establish the first es-fMRI resource in humans and present data from 26 human patients who were undergoing epilepsy monitoring. The dataset comprises pre- and post-surgical whole-brain anatomical (T1-weighted, T1w) and functional (BOLD) MRI imaging at 3T, together with the location of electrodes and the corresponding stimulation protocol parameters. No task was involved, so that the fMRI data can be considered “resting-state” in all cases. Although there are some challenges arising from susceptibility-derived artifacts in the BOLD signal due to the implanted electrode contacts, and some signal artifacts associated with the stimulation, these effects can be analytically reduced and are not extensive enough to impact system-wide inference. As such, this resource and dataset offers unprecedented causal access to network-level organizing principles in the human brain that we expect will grow in the future.

## Methods

### Subjects

26 subjects all gave informed written consent under a research protocol approved by the University of Iowa Institutional Review Board (IRB). Stanford University and Caltech had reliance agreements with the University of Iowa IRB. Written informed consent documents were checked and verified for all participants. See Online-only Table 1 for summary information about stimulation locations.

Participants were patients with medically refractory epilepsy who had elected to undergo neurosurgical treatment for their epilepsy. This is a procedure with a good clinical success rate, provided that seizure foci can be identified. In most patients, scalp EEG together with other clinical information are sufficient to localize the seizure focus. However, a subset of patients have seizure foci that cannot be unambiguously located this way. In this subset of patients, electrodes are implanted into the brain for an extended period so as to provide high-resolution information about the anatomical focus that might be responsible for initiating an epileptic seizure. This approach allows the epileptologist to localize the epileptogenic zone, which in turn can, in many cases, be selectively removed using surgical techniques. In most cases, the patients are also electrically stimulated through some of their electrodes (without fMRI), in order to determine the functional role of brain regions at those locations and guide clinical decisions related to the outcome of neurosurgical resection if it involves those brain regions. These are all common clinical procedures for the treatment of medically refractory epilepsy 17. Patients typically lie in their hospital bed for 1-2 weeks, waiting to have sufficient onset recordings to locate their seizure focus during which they may choose to participate in scientific research.

Our research protocol was conducted when patients consented to participate and when it did not interfere with the clinical protocol. We waited until the seizure focus had been clinically identified and there was no further need to record from the electrodes. Prior to extraction of electrodes and resection for treating epilepsy when deemed appropriate, we introduced our research protocol.

### Safety

Extensive safety testing is described in 15, and relied both on simulations and gel phantom measurements. Potential risks are electrode movement, heating, or current induction as the metal of the electrode, the stimulation current, the time varying magnetic field of the MRI scanner, and the energy deposition from the scanner’s RF coil interact. All of these were found to be well below safety limits and were inspected by the University of Iowa IRB as part of their approval of our protocol.

A Data and Safety monitoring individual from the Department of Neurology at the University of Iowa was responsible for overseeing safety aspects of all of the ongoing studies. An ethics advisory board at the California Institute of Technology also provided initial guidance. Out of all 26 patients, one patient had a seizure during the electrical stimulation protocol; although it remains unclear whether the seizure was in fact triggered by the experimental stimulation. Nonetheless, additional precautionary safety measures were implemented, which included having on-site a trained physician, as well as resuscitation and further monitoring equipment. There were no long-term adverse effects and no complications in this case or any of the other 25 patients.

### Data collection

Pre-electrode-implantation (“pre-op session”; cf. Data Records, below) anatomical MRIs were obtained from all subjects. For these pre-implantation scans we used the following parameters. T1w structural scans were obtained on a 3T GE Discovery 750w (BRAVO, 32 ch head coil, TE = 3.376 ms, TR = 8.588 ms, Flip angle = 12 deg., 1.0 × 1.0 × 0.8 mm voxel size). T2w structural scans were also obtained for some subjects in the same pre-implantation imaging session (CUBE, 32 ch head coil, TE = 86.82 ms, TR = 3200 ms, 1.0x 1.0x 1.0mm voxel size). Up to five eyes-open resting-state BOLD-fMRI sessions were obtained in a subset of subjects as well prior to implantation (4.8min per session, 32 ch head coil, TR = 2260 ms, TE = 30 ms, Flip angle = 80 deg. 3.4 × 3.4 × 4.0 mm voxel size). Some variation in these parameters occurred in the protocol used in the preop session due to scanner availability where a different scanner was used for some subjects. The specifications for each subject can be found in the JSON sidecar information in the data that accompanies each image.

Post-implantation scans (“post-op” session; cf. Data Records, below) included T1w scans for all subjects (MPRAGE, Transmit-receive head coil, TE = 3.44 ms, TR = 1900 ms, TI = 1000 ms, 1.0×1.0×1.0 mm voxel). No T2w scans were obtained, due to safety considerations. Functional scans during the es-fMRI session were obtained on one of two scanners, for earlier datasets on a Siemens 3T TRIO, and later on a 3T Skyra (TR=3000ms; delay in TR=100ms (the electrical stimulation was delivered during this TR delay); TE=30ms, 3 cubic mm isotropic voxels). Stimulation was through an isolated stimulus generator using a biphasic, charge-balanced stimulation delivered across two of the adjacent contacts of a depth electrode (AD-TECH medical instrument, Oak Creek, WI, USA)(Figure 1), as described in further detail in 15. No behavioral or experiential effects were evoked with our brief stimulation, and so the es-fMRI session can be considered cognitively “resting-state.”

**Figure 1:**
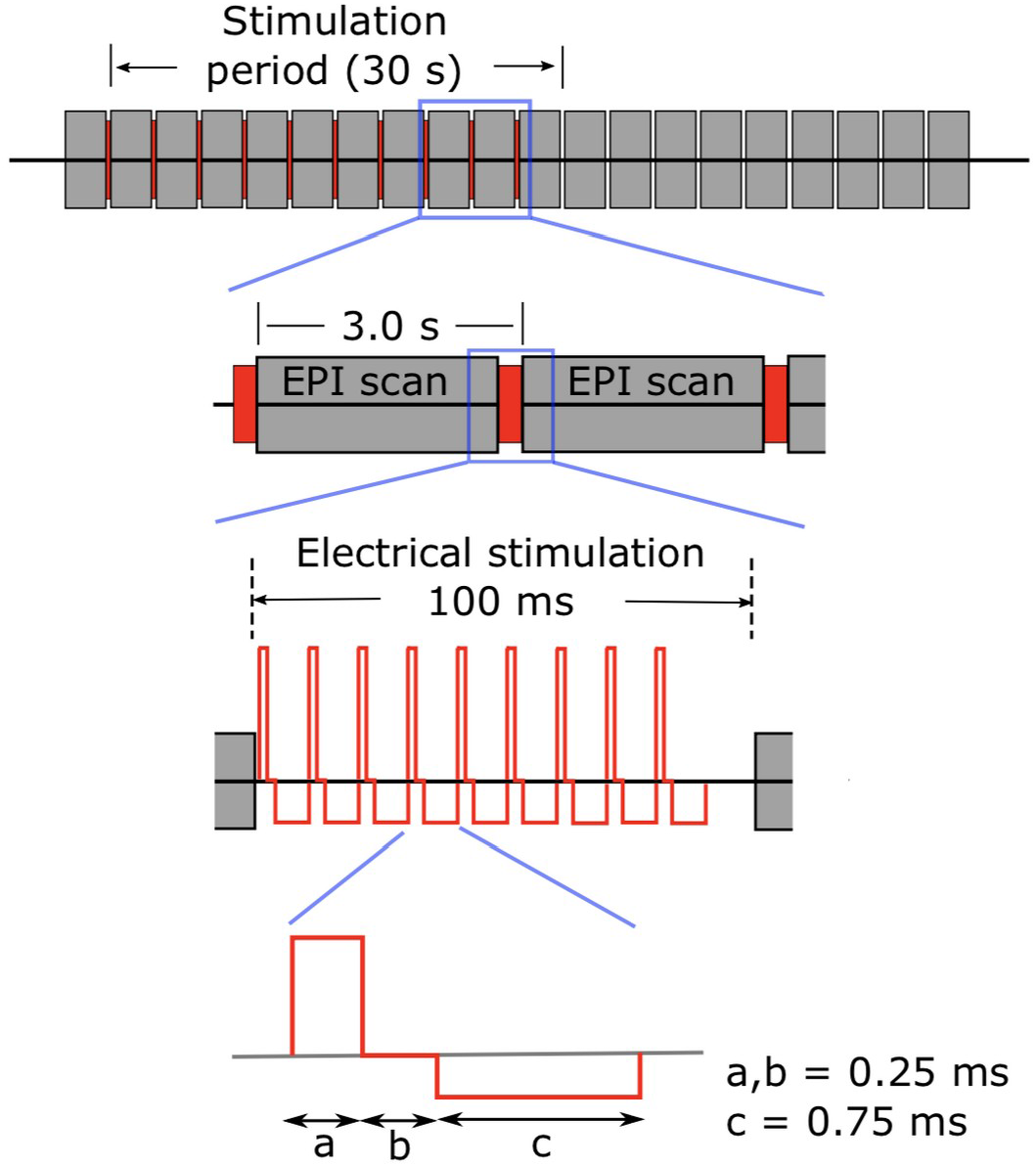
Stimulation paradigm. Electrical stimulation paradigm used for the testing and actual human experiments. Electrical stimuli were delivered so as to be interleaved between EPI volume acquisition, during a 100 ms blank period (no scanner RF or gradient switching). Modified charge-balanced constant-current bi-phasic pulses were used (5-9 pulses at 100Hz; 9-12mA). Stimulation was blocked (ca. 30 seconds ON and OFF) with a total run duration of about 10 minutes (details vary somewhat between subjects). Reproduced with permission from 15.

### Stimulating electrodes: contact co-ordinates and anatomical labels

This is a consolidated spreadsheet that has stimulation contact information for each subject’s es-fMRI runs. For each functional run, information is provided about the stimulation parameters used (i.e. amplitude and duration), the electrode contact anatomical labels (in DTK and Destrieux coordinates), the electrode contact standard coordinates (with respect to two versions of the MNI template), and the hemisphere that was stimulated. The electrode contact-pairs are listed for each es-fMRI run in chA-chB format and the first contact (i.e. chA) gets the leading positive phase of the stimulation. In some cases, two distant sites were stimulated simultaneously. This information is supplied in the ieeg subdirectory in the BIDS file structure (see below).

#### Preprocessed data

The dataset is released with data preprocessed with fMRIPrep 1.5.1rc1 (18; 19; RRID:SCR_016216), which is based on Nipype 1.2.0 (20; 21; RRID:SCR_002502).

##### Anatomical data preprocessing

The preop T1-weighted (T1w) image was corrected for intensity non-uniformity (INU) with N4BiasFieldCorrection 22, distributed with ANTs 2.2.0 (23, RRID:SCR_004757), and used as T1w-reference throughout the workflow. The T1w-reference was then skull-stripped with a Nipype implementation of the antsBrainExtraction.sh workflow (from ANTs), using OASIS30ANTs as the target template. Brain tissue segmentation of cerebrospinal fluid (CSF), white-matter (WM) and gray-matter (GM) was performed on the brain-extracted T1w using fast (FSL 5.0.9, RRID:SCR_002823, 24). Brain surfaces were reconstructed using recon-all (FreeSurfer 6.0.1, RRID:SCR_001847, 25), and the brain mask estimated previously was refined with a custom variation of the method to reconcile ANTs-derived and FreeSurfer-derived segmentations of the cortical gray-matter of Mindboggle (RRID:SCR_002438, 26). Volume-based spatial normalization to MNI152NLin2009cAsym was performed through nonlinear registration with antsRegistration (ANTs 2.2.0), using brain-extracted versions of both T1w reference and the T1w template. The following template was selected for spatial normalization: ICBM 152 Nonlinear Asymmetrical template version 2009c [27,RRID:scr_008796; templateFlow ID: MNI152NLin2009cAsym]

##### Functional data preprocessing

For each of the BOLD runs, the following preprocessing was performed. First, a reference volume and its skull-stripped version were generated using a custom workflow of fMRIPrep. A deformation field to correct for susceptibility distortions was estimated based on fMRIPrep’s fieldmap-less approach. The deformation field was obtained from co-registering the BOLD reference to the same-subject T1w-reference with its intensity inverted 28,29. Registration was performed with antsRegistration (ANTs 2.2.0), and the process regularized by constraining deformation to be nonzero only along the phase-encoding direction, and modulated with an average fieldmap template 30. Based on the estimated susceptibility distortion, an unwarped BOLD reference was calculated for a more accurate co-registration with the anatomical reference.

The BOLD reference was then co-registered to the T1w reference using bbregister (FreeSurfer) which implements boundary-based registration 31. Co-registration was configured with six degrees of freedom to account for distortions remaining in the BOLD reference. Head-motion parameters with respect to the BOLD reference (transformation matrices, and six corresponding rotation and translation parameters) are estimated before any spatiotemporal filtering using mcflirt (FSL 5.0.9, 32). BOLD runs were slice-time corrected using 3dTshift from AFNI 20160207 (33, RRID:SCR_005927). The BOLD time-series were resampled to surfaces on the following spaces: fsnative, fsaverage5. The BOLD time-series (including slice-timing correction when applied) were resampled onto their original, native space by applying a single, composite transform to correct for head-motion and susceptibility distortions. These resampled BOLD time-series will be referred to as preprocessed BOLD in original space, or just preprocessed BOLD. The BOLD time-series were resampled into standard space, generating the following spatially normalized, preprocessed BOLD runs: MNI152NLin2009cAsym. First, a reference volume and its skull-stripped version were generated using a custom workflow of fMRIPrep.

Several confounding time-series were calculated based on the preprocessed BOLD: framewise displacement (FD), DVARS and three region-wise global signals. FD and DVARS were calculated for each functional run, both using their implementations in Nipype (following the definitions by 34). The three global signals were extracted within the CSF, the WM, and the whole-brain masks. Additionally, a set of physiological regressors were extracted to allow for component-based noise correction (CompCor, 35). Principal components were estimated after high-pass filtering the preprocessed BOLD time-series (using a discrete cosine filter with 128s cut-off) for the two CompCor variants: temporal (tCompCor) and anatomical (aCompCor). For aCompCor, components were calculated within the intersection of the aforementioned mask and the union of CSF and WM masks calculated in T1w space, after their projection to the native space of each functional run (using the inverse BOLD-to-T1w transformation). Components were also calculated separately within the WM and CSF masks. For each CompCor decomposition, the k components with the largest singular values were retained, such that the retained components’ time series were sufficient to explain 50 percent of variance across the nuisance mask (CSF, WM, combined, or temporal). The remaining components were dropped from consideration. The head-motion estimates calculated in the correction step were also placed within the corresponding confounds file.

The confound time series derived from head motion estimates and global signals were expanded with the inclusion of temporal derivatives and quadratic terms for each 36. Frames that exceeded a threshold of 0.5 mm FD or 1.5 standardised DVARS were annotated as motion outliers. All resampling was performed with a single interpolation step by composing all the pertinent transformations (i.e. head-motion transform matrices, susceptibility distortion correction, and co-registrations to anatomical and output spaces). Gridded (volumetric) resamplings were performed using antsApplyTransforms (ANTs), configured with Lanczos interpolation to minimize the smoothing effects of other kernels 37. Non-gridded (surface) resampling was performed using mri_vol2surf (FreeSurfer).

#### Example voxelwise GLM analysis

To illustrate an example of the simplest kinds of analyses that could be performed on our data, we conducted a basic GLM analysis. After the minimal preprocessing from fMRIPrep, we applied the following additional preprocessing steps. First, the functional data was spatially smoothed with a Gaussian kernel (FWHM = 8 mm), the data were de-trended, and framewise censoring was applied to volumes with framewise displacement > 0.5mm.

To model the effect of stimulation, we did not use a canonical hemodynamic response function (HRF) and instead estimated the HRF by convolving a boxcar function denoting the electrical stimulation (50-90ms, depending on the actual duration of the stimulation) with an optimized basis set function generated using FLOBS (FSL Linear Optimal Basis Sets)38. This optimization helped account for any non-canonical HRF shapes caused by the stimulation.

A mass univariate analysis was performed for one subject using a linear regression model. The analysis was performed in subject space. Statistical parametric maps shown below were thresholded at an uncorrected voxelwise p < 0.001. The following confounds were used in the regression model: 6 motion parameters, one global signal regressor, anatomical noise components (aCompCor, n=6).

## Data Records

The following data are available and will be periodically updated on the OpenNeuro data sharing platform (https://Openneuro.org;39). The dataset follows the Brain Imaging Data Structure (BIDS;40) version 1.2.2. BIDS organizes the imaging data using a simple folder structure with nested files, each with standardized filenames as per the type of scan. For each scan, the acquired data is converted from DICOM to NIFTI format, and the metadata is stored in an accompanying JSON file. Subsequently, facial features are removed using pydeface to comply with anonymization regulations. Storing data in BIDS format also simplifies the quality testing and preprocessing of the data by making it compatible with BIDS-Apps41 such as MRIQC (for quality assessment;42) and fMRIPrep (for minimal fMRI preprocessing;18). Data is organized into two sessions (namely, “preop” and “postop”, Figure 2).

**Figure 2.**
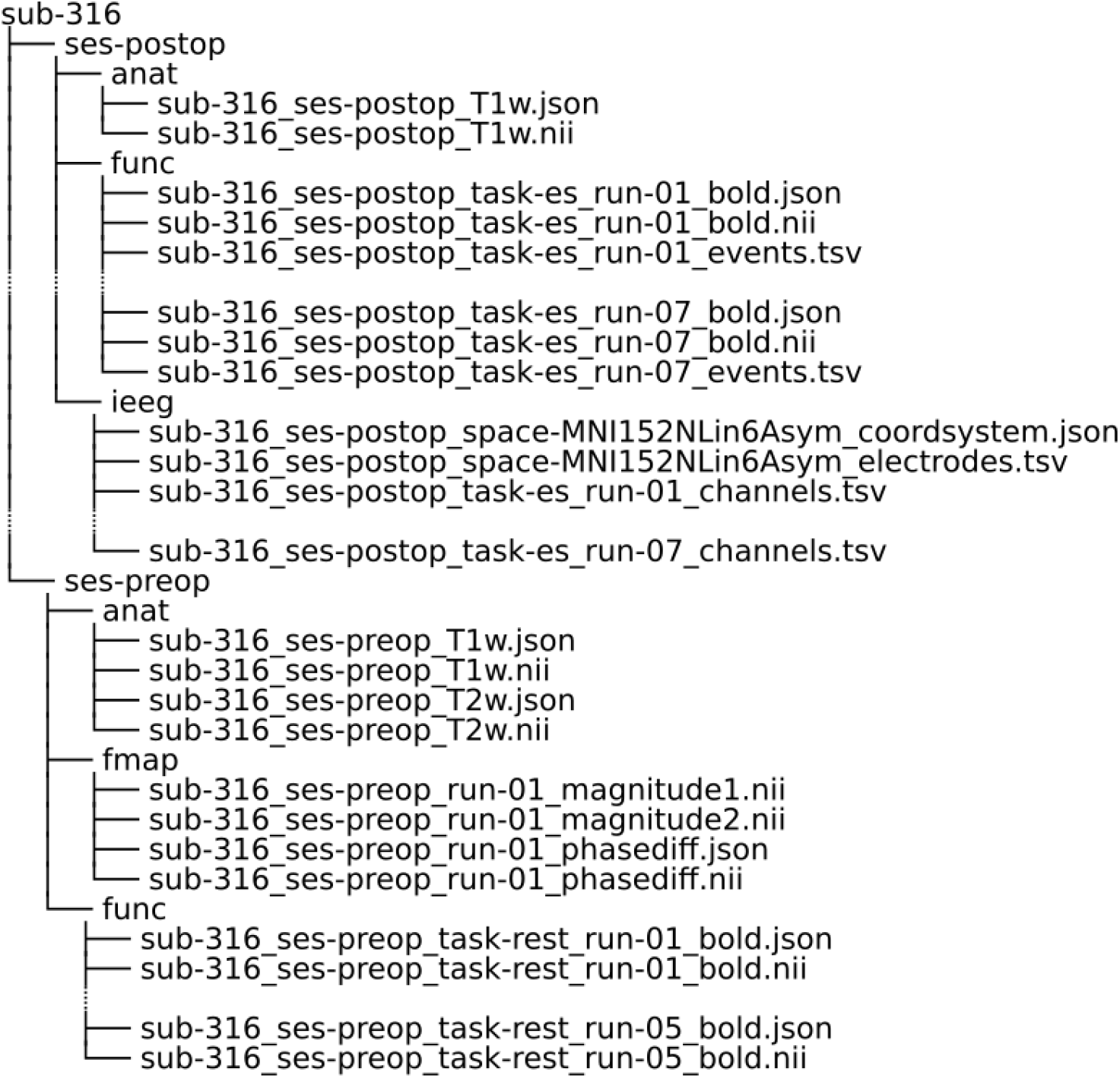
Example of the BIDS data structure for one subject. Here we see the data for subject 316 organized into two sessions, preop and postop. Within the preop there is anatomical, functional and fieldmap data. Within the postop session there is anatomical, functional and electrode information. While the data structure is consistent across subjects, there is some variation regarding data available. For example, the number of postop functional runs can vary between subjects and some subjects do not have preop functional data.

### Pre-op Session

The preop session consists of a pre-implantation MRI protocol including T1w, BOLD fMRI and, in a subset of the patients, GRE phase-difference field maps.

- The pre-op structural images are T1w MRI scans obtained prior to any neurosurgery, which are typically of high quality and are used for registration with post-op images during analysis. Some subjects also have pre-op T2w MRI scans available, to further aid registration and segmentation during analysis.
- The pre-op functional scans are resting-state BOLD fMRI data obtained prior to electrode implantation.
- Pre-op fieldmap images were collected for a subset of the subjects and can be used for field susceptibility distortion correction.

### Post-op Sessions

The postop sessions consist of post-implantation structural MRI and functional MR data as well as associated fieldmaps.

- The post-op structural MRI data are available for all subjects; these show the locations of the stimulating electrodes, as well as of other depth and surface electrodes that have been implanted for monitoring epilepsy.
- The post-op functional data are BOLD fMRI data obtained during concurrent electrical stimulation. These (es-fMRI scans) were obtained post electrode implantation and have blocked electrical stimulation ON or OFF. There was no task, so these functional data are cognitively “resting-state” fMRI.
- Post-op fieldmaps were collected for a subset of subjects to help correct for field susceptibility distortion in the es-fMRI runs.

## Data Quality and Technical Validation

There are a number of potential issues with the es-fMRI technique regarding data quality. In this section we quantify: (1) the quality of the data through quality control metrics, (2) that established preprocessing tools can be applied to the data, (3) that the electrical stimulation did not induce additional movement, (4) validation of the method by showing changes in the BOLD response following electrical stimulation.

### Assessing the quality of es-fMRI data

Using the curated BOLD metrics provided in the crowdsourced MRIQC dataset snapshot43 we compared how the preop resting-state fMRI and the es-fMRI quality-control metrics compared to the broader fMRI dataset quality (Figure 3). This allowed us to contrast both how es-fMRI differs when compared to the preop resting-state scans from the same patients, and how they both relate to a broader overall fMRI quality as obtained in healthy participants typically. MRIQC was performed with the default settings and performed on the images including the noise artifacts (i.e. these have not been masked out).

**Figure 3:**
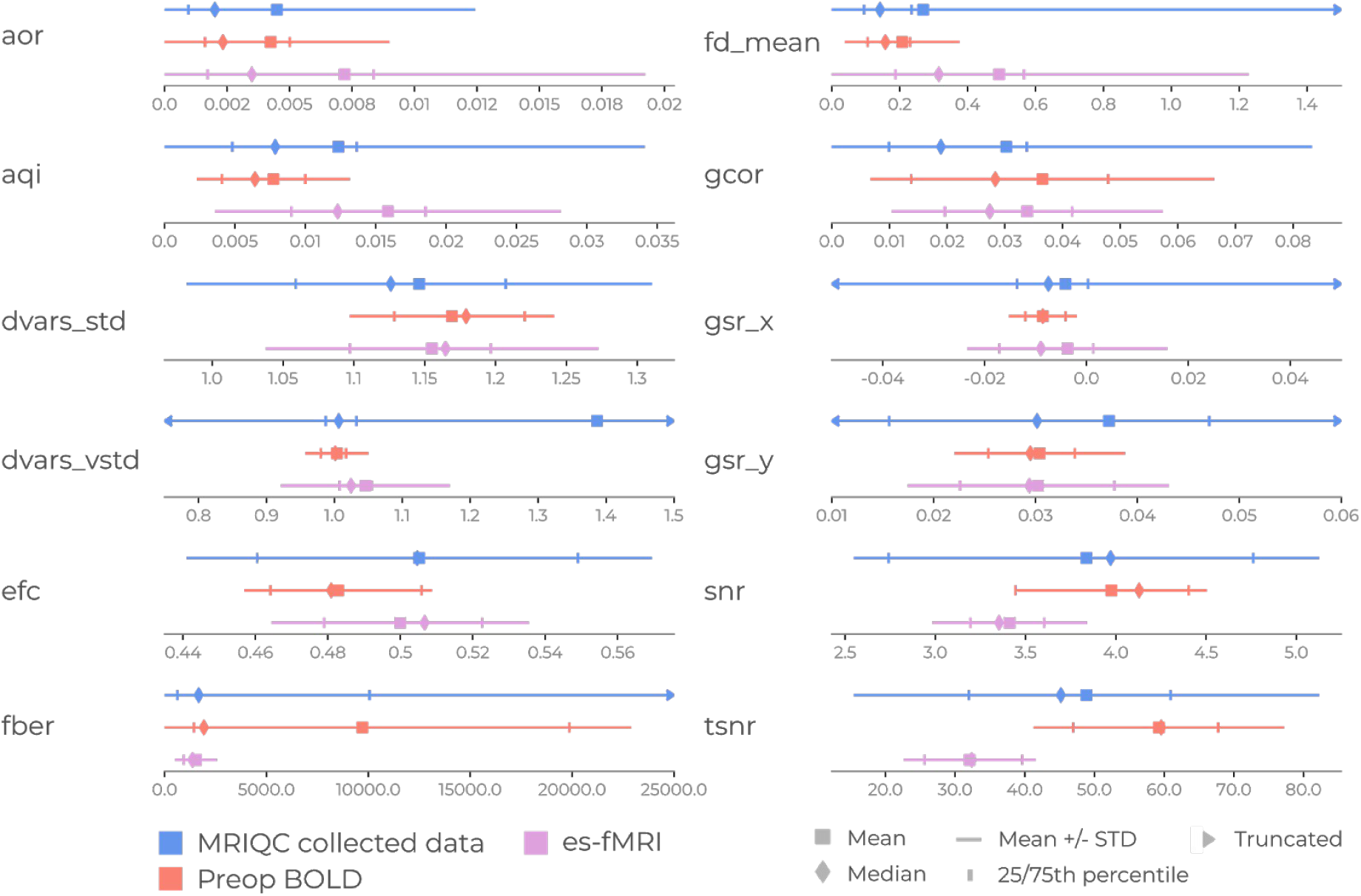
Summary statistics of quality metrics. We compared the preop resting-state BOLD and es-fMRI data with the MRIQC dataset containing the fMRI quality from many datasets. Quality control metrics: *aor* (AFNI outlier ratio), *aqi* (AFNI quality index); *dvars_std* (Standardized derivative of the root-mean-square variance); *dvars_vstd* (Voxelwise standardized derivative of the root-mean-square variance); *efc* (entropy-focus criterion), *fber* (foreground-background energy ratio), *fd_mean* (framewise displacement mean); *gcor* (global correction); *gsr_x* (ghost-to-signal ratio along x-axis); *gsr_y* (ghost-to-signal ratio along y-axis); *snr* (signal-to-noise ratio); tsnr (temporal signal-to-noise ratio)

There are some clear observations in Figure 3. First, the es-fMRI has a loss of signal quality compared to the preop (e.g. SNR, TSNR, AQI). This dropoff is to be expected considering the noise and signal loss due to susceptibility dropout. However, the mean of the es-fMRI’s TSNR falls near the 25th percentile of the MRIQC dataset, showing that the noise levels are not too uncommon for fMRI datasets. Additionally, there is increased movement (i.e. framewise displacement) in the es-fMRI data compared to both the MRIQC dataset and the preop dataset. This observation is evident from the mean framewise displacement for the es-fMRI data, which is above the 75th percentile of both the other two distributions, emphasizing that attention to motion confounds will be critical when analyzing the es-fMRI dataset (see “Usage,” below).

### Preprocessing of es-fMRI data with fMRIPrep

The implantation of electrodes and their magnetic properties induce discontinuities in the B0 field, which translate into substantial distortion artefacts and signal drop-out around the electrodes in the functional data. These distortions accumulate with the inherent distortions typically found around the ventromedial prefrontal area and nearby ear canals in EPI (echo-planar imaging) fMRI. This additional distortion may not be fully corrected with established signal processing routines applied to this data, because of the increased difficulty of image alignment in the presence of substantial distortions. Even if the preprocessing successfully reaches the end of the workflow, there is also the possibility that its quality may be insufficient for reliable analysis. We evaluated these considerations when further preprocessing the data with fMRIPrep.

Data were preprocessed to termination for all subjects, although rejecting some subjects from further analysis may be appropriate after evaluating the quality of the preprocessing. While there were concerns that co-registration between the functional and anatomical data would be poor, this was not the case. We found that the postop functional was co-registered to the preop anatomical scans successfully (Figure 4). Note that we only used the preop T1w images as the anatomical reference within preprocessing of both pre- and postop fMRI, despite the postop T1w being available.

**Figure 4:**
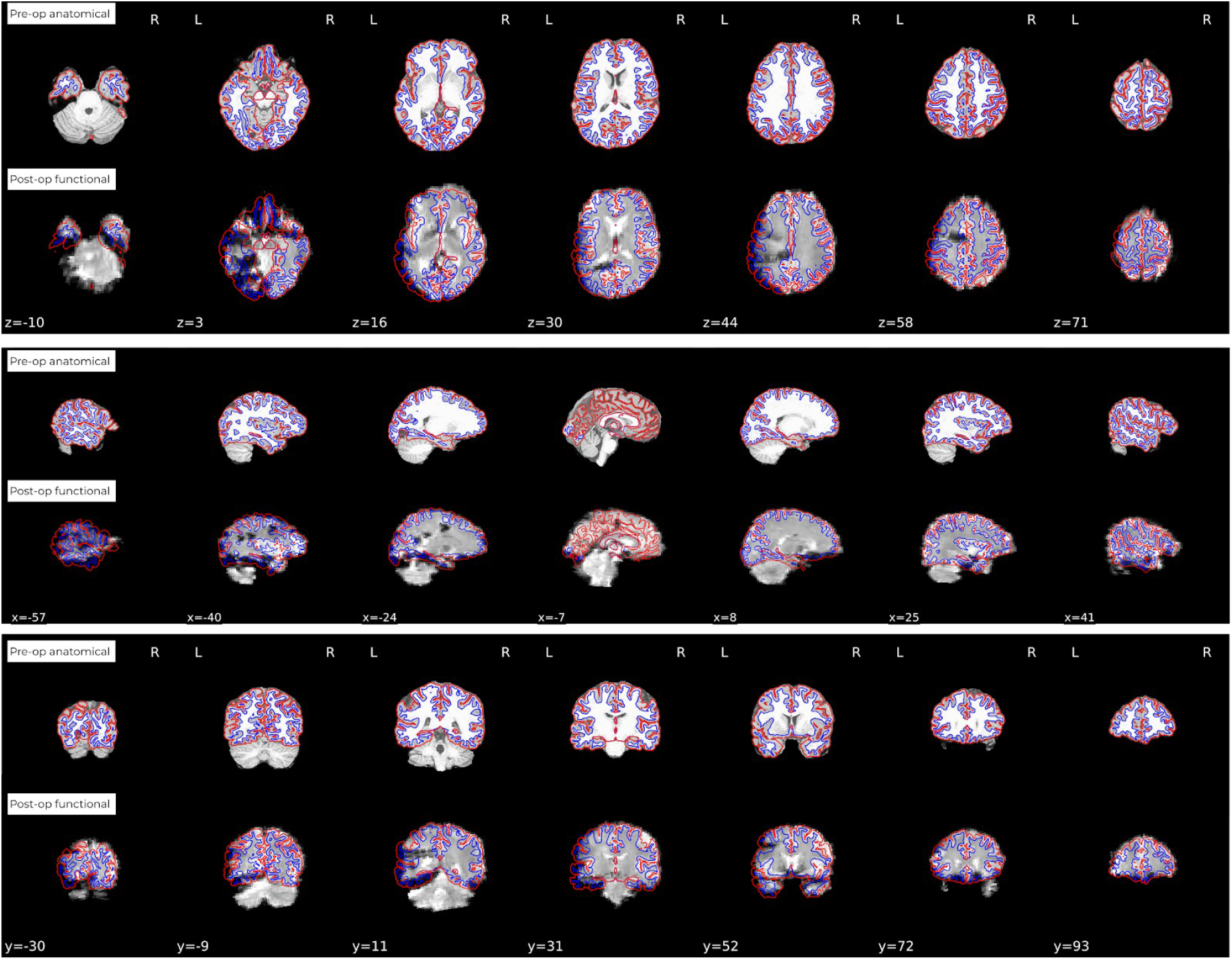
Example of fMRIPrep report of co-registration of postop functional and preop structural images shows extent of noise and distortion on es-fMRI images. Here we can observe that large portions of the left hemisphere of the functional data contains substantial artifacts. However fMRIPrep still co-registers the images adequately despite the distortion. Users of this dataset are strongly encouraged to carefully check each individual subject’s data and co-registration before proceeding to use it in analyses.

Figure 4 illustrates the amount of distortion that exists due to the electrodes in the postop functional data. In this example, considerable noise can be observed in the left hemisphere where the electrodes are implanted. Such noise exists for all subjects to varying extents and, for many types of analyses, should be accounted for (See “usage notes”)

### Movement and global signal as a function of stimulation

Given the often considerable motion, measured by framewise displacement during the entire es-fMRI session, an important quality control check for this dataset is to test whether there is any change in data quality *during* the manipulation (i.e. electrical stimulation ON versus OFF. We were not able to detect any systematic difference across subjects for either the framewise displacement or global signal during stimulation-ON versus stimulation-OFF in the postop functional recordings (Figure 5). Certain subjects have skewed distributions in the framewise displacement leaning towards more movement for either es-ON or es-OFF but this highlights moments of high movement that appear to be independent of the electrical stimulation, as no observable pattern was seen across subjects.

**Figure 5:**
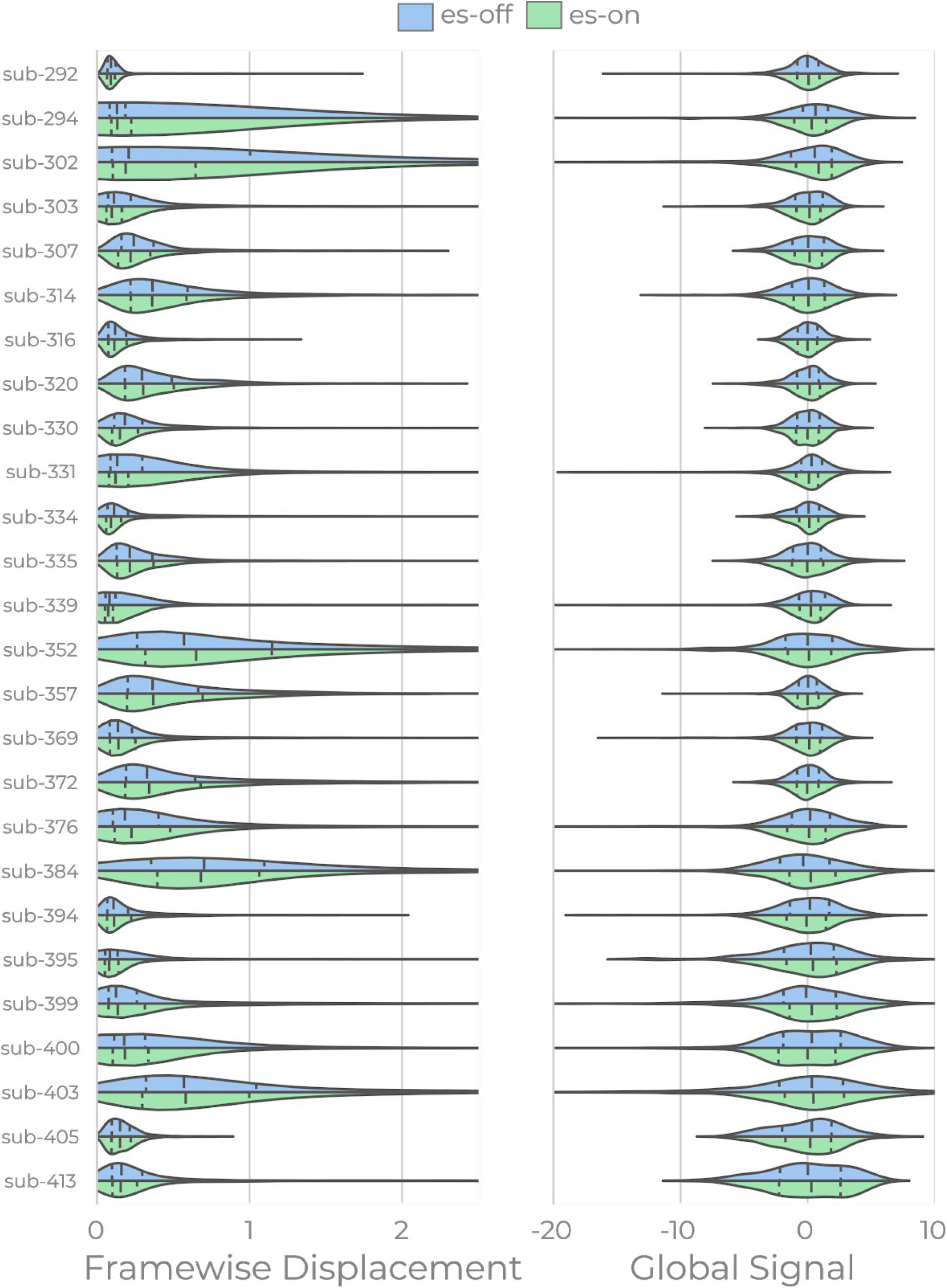
Distributions of framewise displacement (left) and global signal (right) per subject when electrical stimulation was on (green) and off (blue). Framewise displacement is shown on a logarithmic axis. Lines within distribution mark mean an*d standard deviation.*

### Whole Brain Voxel-wise GLM analysis

For one subject, we performed a voxelwise GLM in order to demonstrate that there was change in BOLD relative to the stimulation site. The stimulation site was the left amygdala (Figure 6A). This produced increases in activity both around the amygdala stimulation site but also distributed across the cortex (Figure 6B, p<0.001 uncorrected). Next, we placed a 3mm ROI around the voxel with the peak intensity (crosshair in Figure 6B). The time series of this ROI shows increases in BOLD signal amplitude during stimulation periods that then decrease once the stimulation gets turned off (Figure 6C). Together, these results show: (1) that there is distributed local and distal activation associated with the stimulation; and (2) this increase in activation can be reliably observed in the time series during the stimulation epochs.

**Figure 6:**
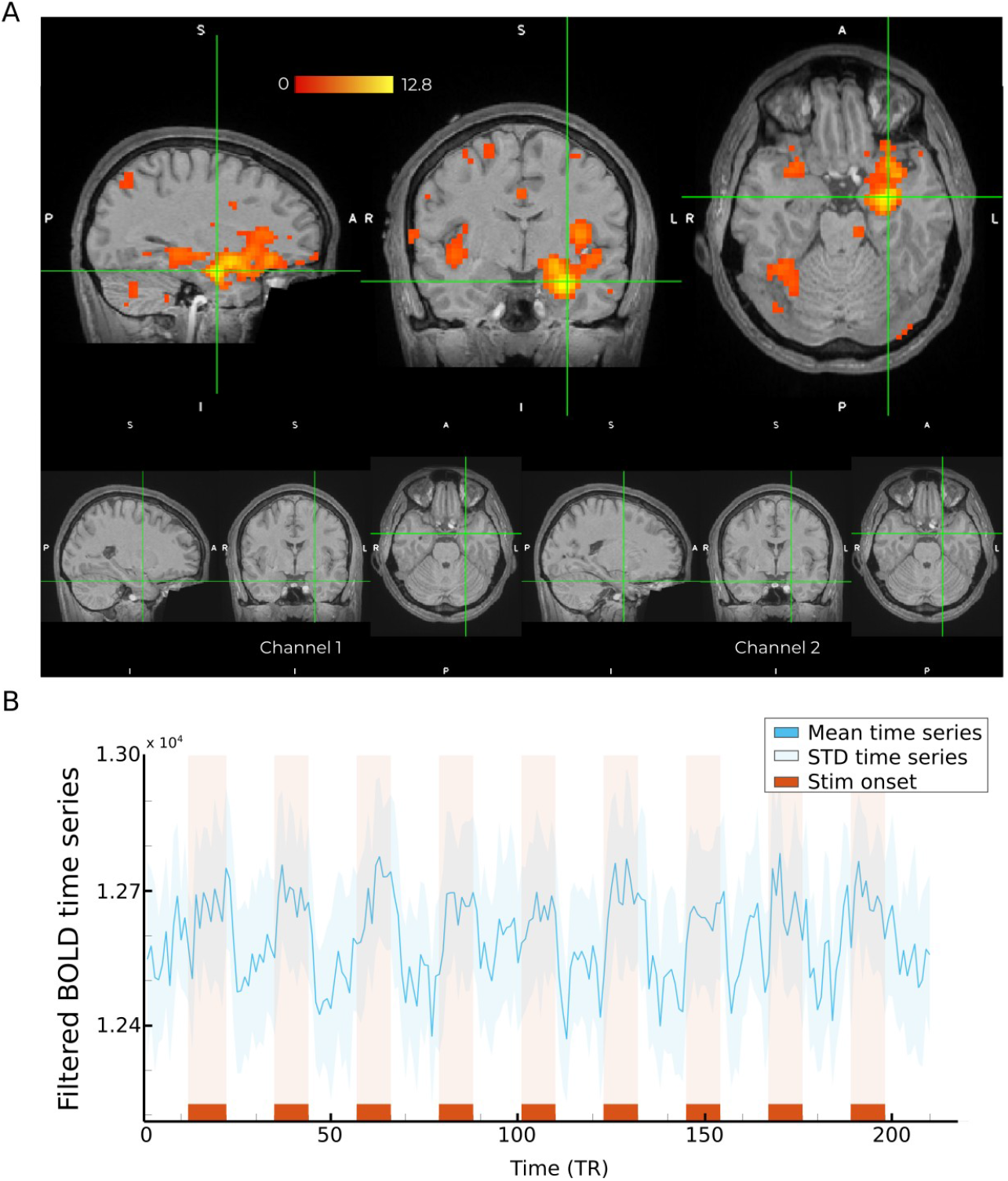
Results of GLM analysis on a single subject (307) during an es-fMRI run with left amygdala stimulation. (A) The top row shows activation in a cluster with the crosshairs centered at the peak intensity voxel (thresholded at p < 0.001). The bottom row shows the location of stimulated electrode (from channel 1 to channel 2). (B) The averaged time course of the filtered BOLD time series (not GLM model fitted response) within a spherical ROI of 3mm centered around the voxel of max intensity shown in the panel. Shaded error bands represent the standard deviation and red bars on the x-axis indicate the electrical stimuli ON periods.

## Limitations and general usage notes

Human es-fMRI is similar to es-fMRI in primate models and at least conceptually similar to optogenetic-fMRI 44,45, and has some notable strengths. First and foremost, it is a unique opportunity to map out functional connectivity in the human brain, and to make inferences about effective connectivity. While resting-state fMRI data is commonly used to derive functional brain networks, its fundamentally correlational nature limits the strength of causal inferences that can be brought to bear on those data. es-fMRI provides a direct causal intervention, and, given certain assumptions, can permit stronger inferences about the causal mechanisms that ultimately generate the data observed 15,16. Secondly, es-fMRI provides valuable information about the network effects of focal DBS, guiding strategies for using DBS to treat psychiatric and neurological disorders. It should be noted in this context that several of the sites that were stimulated in our database are in fact candidates for treating depression. Third, a major benefit of using whole-brain fMRI as the readout in es-fMRI is that it has a whole-brain field-of-view. While the low spatiotemporal resolution of BOLD-fMRI in relation to single neuron studies in animal models remains an important limitation, it is possible and useful to derive information about systems-level brain networks at a mesoscopic scale.

Despite the above strengths, it is also important to discuss the several necessary limitations of es-fMRI and of the present dataset. First, es-fMRI cannot be performed in healthy individuals, and applications thus far have been limited to patients with DBS for treating movement disorders 46–48, or to patients with chronically implanted electrodes for monitoring epilepsy prior to neurosurgery 49 (our case in this paper). Secondly, the electrode placement is determined by clinical criteria; in the case of epilepsy monitoring, electrode locations are fixed and cannot be moved. This results in the relative oversampling of the few locations that are typically targeted in the respective disorders (in our case, medial temporal lobe and medial frontal cortex); on the other hand, it has the benefit of accruing data across subjects who have stimulation in anatomically overlapping regions. Third, depending on the details of the electrode design and the stimulation parameters, electrical stimulation can result in substantial current spread (at least ~3mm, 50), and in any case it is not selective for particular cell types and includes stimulated fibers of passage, unlike optogenetics. Fourth, the implanted electrodes typically introduce susceptibility-derived artifacts (including geometrical distortion and signal loss) on the BOLD-fMRI scan, as we noted earlier. Consequently, those regions are typically masked out of the analysis.

There is yet another potential caveat, which is that, given the low temporal resolution of fMRI, electrical stimulation-fMRI cannot simply be used to infer direct connections between brain regions, as is better gleaned with electrical stimulation - electrophysiology (electrical tract tracing; 51). Instead, the distal BOLD signals produced by focal electrical stimulation should be thought of as network-level effects. There is also the potential confound that electrical stimulation induces conscious precepts of various kinds, leading to indirect changes in BOLD signal as a consequence of cognitive effects. However, this is minimized in our dataset since all patients were at chance in discriminating whether they were electrically stimulated or not; no experiences were evoked. This is likely due to the very brief duration of the stimulation in our protocol. In addition, there are now novel analytic approaches that can be used to accurately infer causal interactions between regions in BOLD data52–54, suggesting that future analyses may be conducted on these data in order to uncover temporal information flow in the human brain.

To deal with the limitations inherent to BOLD-fMRI data, we note some considerations regarding both preprocessing and analysis of the data for those wishing to use this dataset. When preprocessing the raw data, it is recommended to use the preop session’s anatomical images to co-register the postop sessions’s functional data. This is due to the fact that the pre-op anatomical image does not contain as much noise as the post-op counterpart. Analyses using the dataset should consider the additional noise created by the implanted electrodes and electrical stimulation for some analysis strategies. This entails that some strategies should be considered for handling this noise in order to prevent spurious inferences. First, a broad strategy to reduce the noise is to only consider the hemisphere that is contralateral to the stimulation site. This approach could still contain some noise and discard voxels with signal in the ipsilateral hemisphere. Second, masks of “good voxels” per subject can be created in an analysis that spans both hemispheres, by identifying profiles. Work on the second strategy is currently ongoing and aims to be released at a later date. Finally, as noted above, there is considerably more movement in the es-fMRI data than in the pre-op fMRI data, requiring rigorous censoring of time-points and exclusion of runs with high movement; these steps are critical to consider in the analysis of this dataset (e.g. if the average framewise displacement of a run is greater than 0.5, remove the run).

Overall, this novel dataset obtained with es-fMRI in the human brain provides an opportunity to combine causal experimental approaches with whole-brain imaging, which in turn will afford unprecedented opportunities for measuring effective connectivity in the human brain.

## Acknowledgements

Funded by NIH grant U01NS103780 (RP, RA and MH), The Simons Foundation Collaboration on the Global Brain (RA), Knut and Alice Wallenberg Foundation grant 2016.0473 (WHT) and UK Wellcome Trust (CIP) and European Research Council (CIP). The OpenNeuro repository is funded by NIH Grant R24MH117179 (RP).

## Author contributions

Conception and design of the work: HO, MH, RA, CIP

Data collection: HO, MH

Data curation: HO, RN

Data preprocessing, analysis, verification and/or quality control: WHT, RN, OE

Writing (initial draft): WHT, RN, RA

Writing (revising draft): WHT, RN, OE, CIP, JMS, RA

