## Supplementary material for "Human es-fMRI Resource: Concurrent deep-brain stimulation and whole-brain functional MRI": Online Table 1

| Subject ID | Number of Runs | Stimulation Electrode Group | Stimulated Hemisphere |
|---|---|---|---|
| 292 | 3 | Posterior medial frontal depth | Left |
| | 2 | Heschls gyrus depth | Left |
| 294 | 2 | Amygdala depth | Right |
| 302 | 1 | Heschls gyrus depth | Left |
| | 1 | Amygdala depth | Left |
| 303 | 5 | Amygdala depth | Right |
| 307 | 5 | Heschls gyrus depth | Left |
| | 2 | Amygdala depth | Left |
| 314 | 1 | Middle cingulate depth | Left |
| | 1 | Heschls gyrus depth | Left |
| | 1 | Temporal grid | Left |
| | 1 | Amygdala depth | Left |
| | 1 | Anterior cingulate depth | Left |
| 316 | 3 | Heschls gyrus depth | Right |
| | 3 | Amygdala depth | Right |
| | 1 | Posterior hippocampal depth | Right |
| 320 | 2 | Frontal grid | Right |
| | 2 | Heschls gyrus depth | Right |
| | 2 | Amygdala depth | Left |
| 322 | 1 | Anterior medial frontal depth | Right |
| | 3 | Amygdala depth | Right |
| | 2 | Lateral frontal grid | Right |
| 330 | 1 | Parietal grid | Right |
| | 2 | Amygdala depth | Right |
| | 2 | Planum temporal depth | Right |
| | 1 | Amygdala depth & Planum temporal depth simultaneously | Right |
| | 1 | Superior posterior occipital depth | Right |
| 331 | 4 | Heschls gyrus depth | Right |
| | 1 | Planum temporale depth | Right |
| | 1 | Posterior hippocampal depth | Right |
| | 1 | Frontal grid | Right |
| | 1 | Amygdala depth | Left |
| | 1 | Amygdala depth | Right |
| | 1 | Amygdala depth | Left and Right simultaneously |
| 334 | 4 | Planum temporale depth | Right |
| | 3 | Amygdala depth | Left |
| | 1 | Amygdala depth | Right |
| | 1 | Posterior hippocampal depth | Right |
| | 1 | Amygdala depth | Left and Right simultaneously |

| Subject ID | Number of Runs | Stimulation Electrode Group | Stimulated Hemisphere |
| --- | --- | --- | --- |
| 335 | 3 | Heschls gyrus depth | Right |
| | 2 | Amygdala depth | Left |
| | 2 | Anterior insula depth | Left |
| | 1 | Posterior insula depth | Left |
| | 1 | Parahippocampal strip | Left |
| | 1 | Temporal grid | Right |
| 352 | 7 | Amygdala depth | Left |
| | 2 | Heschls gyrus depth | Left |
| 357 | 4 | Heschls gyrus depth | Left |
| | 2 | Frontal grid | Left |
| | 1 | Middle medial frontal depth | Left |
| | 1 | Temporal grid | Left |
| | 1 | Posterior medial frontal depth | Left |
| 372 | 4 | Heschls gyrus depth | Left |
| | 3 | Amygdala depth | Left |
| 376 | 1 | Amygdala depth | Right |
| | 2 | Frontal grid | Right |
| | 1 | Posterior insula depth | Right |
| | 1 | Heschls gyrus depth | Right |
| 384 | 4 | Amygdala depth | Right |
| 395 | 3 | Amygdala depth | Left |
| | 1 | Insula depth | Left |
| 399 | 4 | Anterior cingulate genu depth | Right |
| | 2 | Amygdala depth | Right |
| | 2 | Heschls gyrus depth | Right |
| 400 | 2 | Heschls gyrus depth | Left |
| | 1 | Amygdala depth | Left |
| | 1 | Parietal - middle cingulate depth | Left |
| | 1 | Posterior hippocampus depth | Left |
| 403 | 1 | Temporal grid | Left |
| | 1 | Heschls gyrus depth | Left |
| | 3 | Hippocampus depth | Right |
| 405 | 2 | Posterior insula depth | Left |
| | 1 | Amygdala depth | Right |
| | 1 | Amygdala depth | Left |
| | 1 | Hippocampus depth | Right |
| | 1 | Anterior insula orbitofrontal cortex depth | Left |
| 413 | 1 | Frontal operculum ofc depth | Right |
| | 1 | Cingulate genu depth | Right |
| | 1 | Inferior posterior insula depth | Right |
